# Orthogonal Representations of Object Shape and Category in Deep Convolutional Neural Networks and Human Visual Cortex

**DOI:** 10.1101/555193

**Authors:** Astrid A. Zeman, J. Brendan Ritchie, Stefania Bracci, Hans Op de Beeck

**Author notes:** corresponding author: Correspondence; Correspondence Address: Brain and Cognition Department, KULeuven. Tiensestraat 102, Leuven 3000 Belgium.

## Abstract

Deep Convolutional Neural Networks (CNNs) are gaining traction as the benchmark model of visual object recognition, with performance now surpassing humans. While CNNs can accurately assign one image to potentially thousands of categories, network performance could be the result of layers that are tuned to represent the visual shape of objects, rather than object category, since both are often confounded in natural images. Using two stimulus sets that explicitly dissociate shape from category, we correlate these two types of information with each layer of multiple CNNs. We also compare CNN output with fMRI activation along the human visual ventral stream by correlating artificial with biological representations. We find that CNNs encode category information independently from shape, peaking at the final fully connected layer in all tested CNN architectures. Comparing CNNs with fMRI brain data, early visual cortex (V1) and early layers of CNNs encode shape information. Anterior ventral temporal cortex encodes category information, which correlates best with the final layer of CNNs. The interaction between shape and category that is found along the human visual ventral pathway is echoed in multiple deep networks. Our results suggest CNNs represent category information independently from shape, much like the human visual system.

## Introduction

In recent years, the performance of Deep Convolutional Neural Networks (CNNs) has improved significantly, such that they are able to meet^1-3^, and even surpass^4^ human performance in classifying objects. In light of these impressive findings, these artificial networks are increasingly compared to their biological counterparts, resulting in an accumulation of evidence for their use as a benchmark model of visual object recognition^5,6^. For example, the internal representations of CNNs show correspondence with human ventral temporal cortex (VTC) as measured by fMRI, as well as with primate inferotemporal cortex (IT) measured using single cell recordings^7-12^. The correspondence between deep networks and neural representations along the visual pathway has even allowed for accurate neural response prediction of single-cell recordings in IT^9^ as well as fMRI^13^. Representational similarities have been further extended from the spatial into the temporal domain, with results showing a corresponding ordering of processing between CNNs and the human visual brain using MEG^14^. These accumulating findings showcase the ability of CNNs to model neurons from single unit responses to entire populations, spanning the multiple scales and dimensions used to study neural activity, and making CNNs one of the best models to date for studying vision in the human and primate brain.

While these feats are impressive, it is unclear to what extent these results are easily interpretable in terms of category representations. Object category information can often be confounded with low-level visual features, such as colour, texture, and shape^15^. In this paper, we highlight the significant interaction between shape and category that is known to occur in natural images^16^ and address the possibility that these networks may distinguish between object categories by relying upon visual features, such as shape, rather than high-level category representations. Indeed, the shape similarity of objects has been capitalised on in the machine learning field to improve performance^17^. CNNs are proficient at representing the perceived shape of objects, as opposed to their physical shape^18^ and there are claims that CNNs rely heavily upon shape information for classification^19^. Two-dimensional regular vs irregular shape representations have been found in monkey IT, which are highly comparable to late layers of CNNs^12^. Furthermore, CNNs mimic a behavioural bias in humans known as the “shape-bias”, which is the preference to categorise an object based on shape rather than colour^20^. Given that these networks are adept at representing object shape, it is possible they are taking advantage of shape-based features, instead of category information, to classify object images.

Recent neuroimaging studies have begun to de-cofound category from visual features, including shape, in order to investigate their interaction along the visual ventral pathway^10,16,21,22^. VTC in humans is one of the main category-selective areas^23^, distinguishing, for example, between animate and inanimate objects^24,25^. To build up this category-related representation, visual information is processed in a series of stages along the ventral visual pathway, from primary visual cortex (area V1) through to VTC^23^. In recent years, the exact role of VTC has come under question, in particular whether this area encodes category-specific information, or simply the low-level visual properties associated with category, such as colour, shape, size and texture^15,26,27^. Proklova, Kaiser & Peelen^22^ found that VTC encodes texture and outline alongside category-specific information that is not present in earlier visual areas. Another higher visual area, lateral occipitotemporal complex (LOTC), was found to encode category-associated shape properties as well as category-selective information^21^. Other category-orthogonal object properties, including size, position and pose, show higher population decoding performance in monkey IT (analogous to human VTC) compared to early visual areas, contrary to what was previously believed^10^. Indeed, the majority of visual object representations in IT may be accounted for by object shape, or other low-level visual properties, rather than category^28^. Nevertheless, studies that explicitly de-confound category from more low-level properties suggest that the category selectivity cannot be fully explained by these other properties^10,16,21^, and point towards a so-called feature-dependent categorical code^15^.

In this paper, we explicitly dissociate shape from category in two stimulus sets to determine: (i) how CNNs represent object shape and category when they are independent from one another; and (ii) how these artificial representations correspond with shape and category representations in human visual cortex. Using two carefully designed stimulus sets, which orthogonalise shape and category, we assess four top-performing CNNs in their ability to represent category independently from shape layer by layer. Taking the same two stimulus sets, we measure human fMRI responses when viewing these images and assess the interaction between shape and category along the visual ventral stream. Finally, we compare artificial representations with human fMRI responses for the same two stimulus sets, to evaluate how closely CNNs reflect biological representations.

## Methods

We aimed to determine the relationship between models of shape and category, CNNs, and neural responses in the human visual ventral pathway. We tested object shape and category representation in four top-performing CNNs and compared this with behavioural ratings of shape and category as well as human fMRI response patterns from experiments in two previous studies^16,29^. Below we describe participants, stimulus sets, CNN architectures, the neuroimaging experiments, and data analysis.

### Participants

All participants gave written informed consent. All experiments were approved by the Ethics Committee at KU Leuven and the University Hospitals Leuven. All methods were performed in accordance with the relevant guidelines and regulations. For the behavioural ratings, each stimulus set was rated by an independent group of participants (N= 4 for set A; N = 16 for set B). For the neuroimaging experiments, there were 15 participants (8 females, mean age of 30 years) scanned in fMRI experiment A, none whom were excluded. There were also 15 participants (8 females, mean age of 24 years) scanned for fMRI experiment B, with one person who was excluded due to excessive head motion. All subjects had normal or corrected vision.

### Stimulus sets

The stimuli in both experiments were designed to dissociate shape from category information. Both stimulus sets are grayscale images of objects on a white or grey background, centred at the origin and presented at a normal viewing angle (see Figure 1). Set A contains 32 unique images, divided into 2 equally sized categories (animal vs non-animal) and 2 equally sized groups of shapes (low and high aspect ratio). Set B contains 54 images divided into 6 object categories (minerals, animals, fruit/veg, music, sport and tools) and 9 shape types. The model design for each stimulus set, which orthogonalises shape from category, is illustrated in Figure 1. For additional information about the stimulus sets, refer to Ritchie and Op de Beeck^29^ and Bracci and Op de Beeck^16^, for Set A and B respectively.

**Figure 1:**
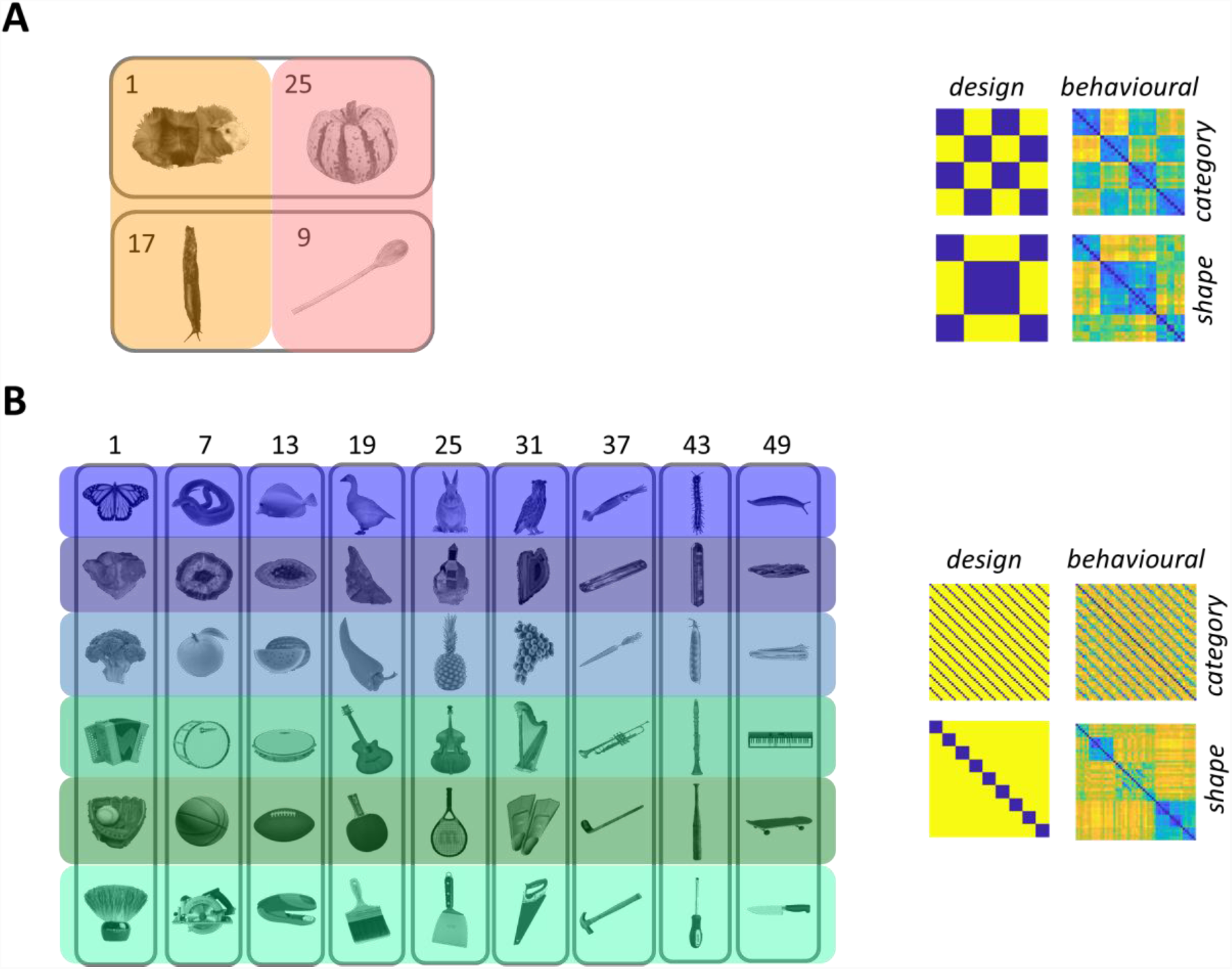
(A) 32 stimuli in 2 categories (animal and non-animal), (B) 54 stimuli in 6 categories (animals, minerals, fruit/vegetables, music, sports equipment, tools). Left: Each category division is highlighted by a distinct colour. Common shape information is circled in grey. Numbers indicate indexing for RDMs. Due to copyright restrictions, not all images are shown in Set A and the ones displayed are representative. Set A images are published in compliance with a CC BY-SA license (https://creativecommons.org/licenses/by-sa/3.0/) and their sources are: guinea pig (https://commons.wikimedia.org/wiki/File:AniarasKelpoKalle.jpg by Tavu); squash (https://commons.wikimedia.org/wiki/File:Festival-Squash.jpg by Evan-Amos); slug (Black Slug at Aggregate Ponds, https://www.flickr.com/photos/brewbooks/2606728819 by brewbooks); and wooden spoon (https://upload.wikimedia.org/wikipedia/commons/7/7b/Wooden_Spoon.jpg by Donovan Govan). Images have been changed to greyscale and have the background removed. The final two images have also been rotated. Set B images are published in compliance with a CC-BY license (https://creativecommons.org/licenses/by/4.0/) and are re-used from Figure 5a in Kubilius, Bracci and Op de Beeck^18^. Right: Shape and category RDMs. The design models are based on the experimental design. The behavioural models are obtained via multiple object arrangement^31^; see methods.

To confirm that shape was not predictive of category information for each of the stimulus sets, we analysed the images using low-level GIST descriptors^30^ and tested how well these visual features predicted shape or category using Linear Discriminant Analysis (LDA). GIST provides a low dimensional representation of an image based on spectral and coarsely localised information. We defined the GIST descriptors to include 8 orientations over 8 scales and combine this with LDA. For Set A, we ran a two-way classification using a leave-one-level out procedure, for example, training on bar stimuli and generalising to blob stimuli to test for animacy classification. For Set B, we followed a six-way classification using a leave-one-level out test procedure, permuting across all possible groups of train and test combinations and averaging across results. For example, we selected six shape clusters of the total nine, trained an LDA on GIST descriptors from five clusters (5×6 = 30 images) and tested whether the algorithm could predict the 6 different categories from the held out images. All six-way shape and category combinations were tested and averaged.

### Behavioural data

Each stimulus set was rated on object category and shape properties by means of the multiple object arrangement method^31^. Participants rated similarity in two task contexts: for *object category*, “arrange the images based on the semantic similarity among objects”; for *object shape*, “arrange the images based on perceived object shape similarity”. These models, based on behavioural data, are meant to better represent the stimulus psychological space relative to the stricter *design-based* models (2 categories x 2 shape types in set A; 6 categories x 9 shape types in set B). For example, in Set B, the *design-based* shape model represents the 9 different shape types as equidistant from one other, whereas the *behaviour-based* shape model is sensitive to further variation between the 9 shape types in terms of between-type similarity. The behaviour-based model for Set B illustrates that elongated objects (the final 3 shape types), regardless of their orientation, are perceived as being more similar to each other relative to round objects (the first 3 shape types), which is not visible in the design-based model. Figure 1A and 1B depicts both *design-based* and *behaviour-based* models.

### fMRI Experiments

Here we provide a summary of the fMRI procedures and analyses, the full details are provided in Ritchie and Op de Beeck^29^ for experiments using Set A and Bracci and Op de Beeck^16^ for Set B.

#### Preprocessing and Analysis

All imaging data was pre-processed and analysed using SPM and MATLAB. For each participant, fMRI data was slice-time corrected, motion corrected (using spatial realignment to the first image), coregistered to each individual’s anatomical scan, segmented and spatially normalised to the standard MNI template. Functional images were resampled to 3 x 3 x 3 mm voxel size and spatially smoothed by convolving with a Gaussian kernel of 6mm FWHM for Set A and 4mm FWHM for Set B^32^. After pre-processing, a GLM was used to model the BOLD signal for each participant, for each stimulus, at each voxel. Regressors for the GLM included each stimulus condition of interest (32 for A, 54 for B) and 6 motion correction parameters (x, y and z coordinates for translation and rotation). Each predictor had its time course modelled as a boxcar function convolved with the canonical haemodynamic response function, producing a single estimate for each voxel per predictor for every run. The beta weights fitted to each GLM were used to create Representational Dissimilarity Matrices (RDMs) for each participant (defined below).

#### Regions of Interest (ROIs)

Neural representational content was investigated in three main ROIs in visual cortex: primary visual cortex (V1), and ventral temporal cortex (VTC), which was split into posterior (VTC post) and anterior (VTC ant) halves. These ROIs were chosen for their relevance in both object shape and category information processing^23^. VTC is bounded laterally by the occipitotemporal sulcus (OTS), posteriorly by the posterior transverse collateral sulcus (ptCoS) and anteriorly by the anterior tip of the mid-fusiform sulcus (MFS)^23^. ROIs were defined at the group level by combining the anatomical criteria above (using the Neuromorphometrics atlas in SPM) with functional criteria (all active voxels for the contrast of all conditions versus baseline that responded to visual information exceeding the statistically uncorrected threshold of p < 0.001 in a second-level analysis). For further details on ROI definition, please refer to Bracci, Kalfas & Op de Beeck^33^ where the exact same ROI criteria were applied. We used a two-factor repeated-measures Analysis of Variance Model (ANOVA) to assess the interaction between two within-participant factors: conditions (shape, category) and area (V1, VTC post and VTC ant).

### Deep Neural Network Architectures

Each architecture consists of multiple convolutional layers followed by pooling operations and fully-connected layers. For each CNN, which was pre-trained on the ImageNet dataset^34^, we ran a forward pass of each image in the stimulus set through the network. We output the activation of weights in each layer, resulting in a matrix with size of the *nodes per layer* times *the stimulus set* (32 for A, 54 for B). We calculated *1 - correlation* for each activation pattern of one stimulus with another to obtain an RDM with size N x N, where N = the number of stimulus conditions (32 x 32 for A, 54 x 54 for B). We did not include final softmax classification layers in our analysis, since we were interested in the structure of layer representations and not classification performance per se.

#### CaffeNet

CaffeNet is an implementation of AlexNet^1^ in the Caffe deep learning framework^35^.CaffeNet is an 8-layer convolutional neural network (CNNs) with five convolutional layers and three fully connected layers.

#### VGG-19

VGG-19^3^ was the top ranking CNN for single object localisation in ILSVRC 2014, and second-running in image classification^34^. VGG-19 consists of 19 weighted layers with an additional softmax read-out layer for classification. The architecture contains 16 convolutional layers separated by five max pooling layers, with the final 3 layers being fully-connected.

#### GoogLeNet

GoogLeNet^2^, also known as InceptionNet, was the top-performing architecture for image classification in ILSVRC 2014^34^. GoogLeNet is a 22-layer deep network, when counting only parameterised layers, or 27 layers deep if including pooling operations. All convolution, reduction and projection layers use rectified linear activation. The bottom layers of the network follow conventional convolutional neural network architecture, consisting of chained convolutional operations followed by max pooling. The top layers of the network replace multiple fully-connected layers with an average pooling layer, a single fully connected layer and a classification layer. The middle layers of the network differ substantially from traditional convolutional neural network structure, consisting of stacked “inception” modules, which are miniature networks containing one max pooling and 3 multi-sized convolution operations (1 x 1, 3 x 3 and 5 x 5 convolutions) in parallel configuration. Convolution operations inside inception modules are optimised with dimensionality reduction, by preceding expensive 3 x 3 and 5 x 5 convolution operations with 1 x 1 convolutions. Inception modules allow for increased width of the network, as well as depth, while maintaining a constant computational budget.

#### ResNet50

ResNets are a family of extremely deep architectures that won the ILSVRC classification task in 2015^36^. ResNet50 contains 50 stacked “residual units”, which use a split-transform-merge strategy to perform identity mappings in parallel to 3×3 convolutions with rectification. ResNets, like GoogLeNet^2^, are multi-branch architectures, containing only 2 branches (performing identity projection and 3×3 convolutions) instead of GoogLeNet’s maximum 4 branch inception modules (performing multi-size convolutions). Identity mappings perform a key role in the architecture’s success, forcing the network to preserve features, rather than learn entirely new representations at every layer, as is the case with conventional CNNs^37^. The final 3 layers of ResNet50 are identical in design to GoogleNet, performing average pooling, transformation to 1000 dimensions using full connections and softmax classification (not included in our analysis).

### Representational Similarity Analysis

We used Representational Similarity Analysis (RSA) to quantitatively compare CNN representations per layer with design models, behavioural ratings, and with fMRI neuroimaging data. RSA compares RDMs, which characterise the representational information in a brain or model^38^. Given a set of activity patterns (biological, behavioural or artificial) for a set of experimental conditions, the dissimilarity between patterns is computed as 1 minus the correlation across the units that compose the patterns. RDMs are symmetrical about a zero diagonal, where 0 denotes perfect correlation. RSA assesses second-order isomorphism, which is the shared similarity in structure between dissimilarity matrices^39^. Spearman rank order correlation was used to compare dissimilarity matrices, since the relationship between RDMs cannot be assumed to be linear^38^. In cases where there was any dependency relationship between shape and category RDMs (visible in the Set A behavioural data), we used partial correlation. We determined the significance of every correlation by comparing it with a null distribution obtained by randomly permuting the RDM labels and then calculating dissimilarity relationships 1000 times.

## Results

### Behavioural Data

For each stimulus set, participants provided similarity judgments for the shape and category dimension (see Figure 1, right column). For Set A, we found a significant correlation between the behavioural models for shape and category (Spearman’s ρ = 0.4753, *p* < 0.001 permutation test with 10000 randomisations of stimulus labels) and so partial correlations when carrying out RSA with Set A behavioural models. For Set A, as expected, behavioural and design category models strongly correlate with one another (ρ = 0.8555, *p* < 0.001) and design shape strongly correlates with behavioural shape (ρ = 0.7849, *p* < 0.001). For Set B, we found no significant correlation between behavioural models for shape and category (ρ = 0.006, *p* = 0.8209). Again, as expected, shape behavioural and design models were significantly correlated (ρ*=* 0.4145, *p* < 0.001) and category behavioural and design models were also significantly correlated (ρ*=* 0.6195, *p* < 0.001).

### Low-level Shape Analysis of Stimuli

Using GIST^30^ descriptors of each image and combining this with LDA, we confirmed that category could not be predicted based upon these low-level descriptors whereas shape could, demonstrating that our stimulus sets were properly orthogonalised. LDA with GIST predicted shape above chance level, at 87.5% for Set A and 69% for Set B. Category was predicted below chance level, at 37.5% for Set A and 10% for Set B.

### Shape and category RSA on all CNN layers for Stimulus Sets A and B

Figure 2 illustrates layer-by-layer RSA between the CNN representations and the shape and category models and behavioural data in the two stimulus sets. Note that all RSA using Set A behavioural models involved partial correlations (explained above in Behavioural data). Looking across all networks, in the first layer of all CNNs, shape is already represented above the significance threshold in most cases, whereas category is not. Shape correlations at the first layer of CNNs are lower and closer to the significance threshold for Set A (design 0.12 < ρ < 0.22, behavioural 0.12 < ρ < 0.24) than Set B (design 0.26 < ρ < 0.44, behavioural 0.24 < ρ < 0.36). CaffeNet shows the highest correlation in shape information at the first layer with both behavioural and design models for both stimulus sets. In CaffeNet, there is a single rise and fall in shape information, except in the Set A behavioural model. In all other networks, shape correlations fluctuate along the layers, with peaks at different layers before decreasing at the final layer in all cases except for Set A GoogLeNet and ResNet50. For Set A, shape correlations remain relatively high at the final layer (design 0.34 < ρ < 0.51, behavioural 0.29 < ρ < 0.59). In contrast, for Set B, shape correlation levels increase in the networks before falling in the final layers of all networks, to below their first layer levels for the design model correlations (0.11 < ρ < 0.14), or to roughly their initial values for the behavioural model correlations (0.32 < ρ < 0.36). For all networks, category information remains low across the majority of layers, hovering at or below the significance level until the final few layers, where it increases above the significance threshold to peak at the final layer. At the final layer, for Set A, category correlations reach between 0.31 < ρ < 0.42 for design models and between 0.34 < ρ < 0.42 for behavioural. For Set B, category correlations reach between 0.11 < ρ < 0.21 for design models and between 0.24 < ρ < 0.37 for design models at the final layer.

**Figure 2:**
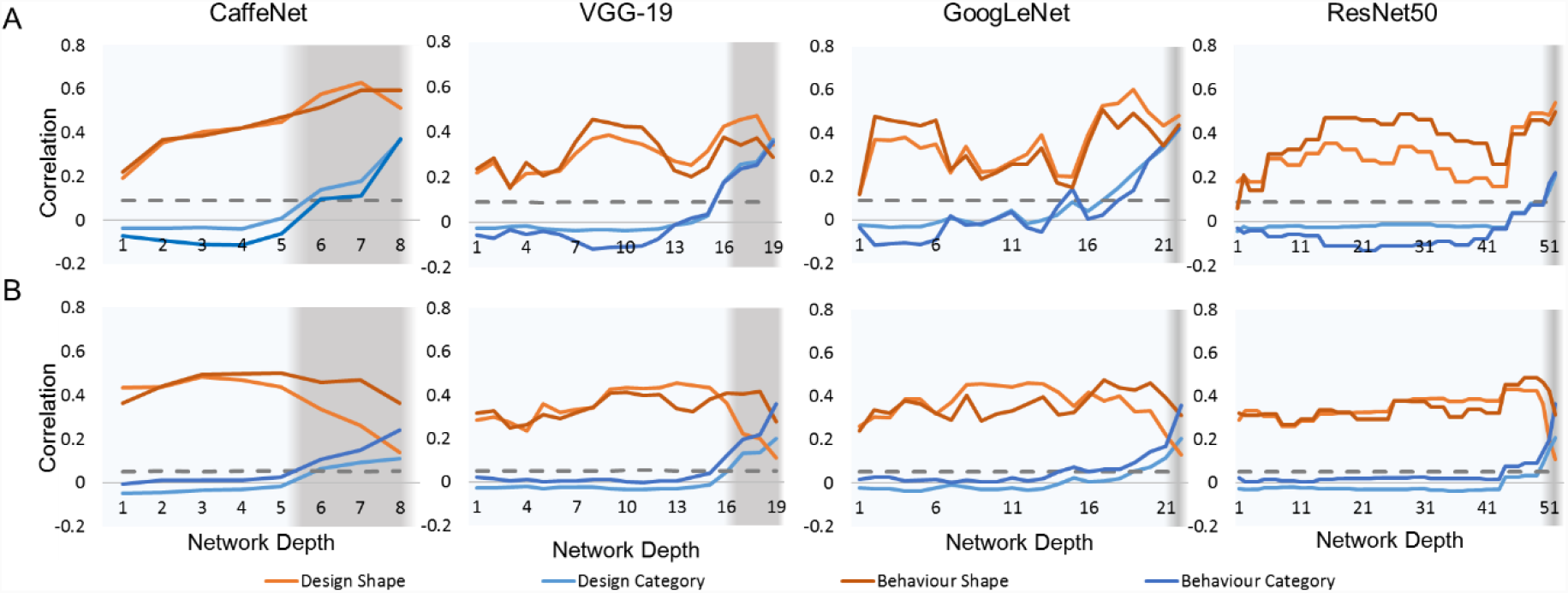
Correlation between layers in CNNs and shape (orange/red) versus category (blue) in Set A (top row) and B (bottom row). The horizontal axis indicates network depth and the vertical axis indicates correlation (Spearman’s ρ). For GoogLeNet and ResNet architectures, the correlations shown are for 3×3 convolutional operations, while other parallel operations (projections and convolutions of different sizes) are omitted. Dashed line indicates significance threshold of p < 0.05. Grey shading indicates fully-connected layers.

To investigate the interaction between shape and category and CNN layers, we tested correlation values in a 2 × 2 ANOVA with Layer (modelled linearly with intercept and slope) and Condition (Shape or Category). *Table 1* summarises the statistical results of the main effects (layer, condition) and their interaction in CNNs and models. For Set A, for both types of models across all networks, layer has a highly significant main effect and condition is also significant (*Table 1*) which suggests that correlation values can be predicted given the CNN layer and the condition of interest (shape or category information). Their interaction is significant in GoogleNet and VGG-19, but not in CaffeNet and ResNet50, suggesting that as category increases, shape decreases significantly in two out of the four networks tested. For Set B, across all networks, condition is highly significant, and layer has a significant main effect in behavioural model correlations, however regarding design model correlations, layer is only significant in one CNN (ResNet50). This suggests that it is possible to make significant predictions of behavioural shape and category judgements given CNN layer information, however this prediction does not extend to design models of shape and category. Condition is highly significant across all networks, and the interaction between layer and condition is significant for both models and CaffeNet, and the design model and GoogleNet.

**Table 1:** 2 × 2 ANOVA results of Layer (modelled linearly with slope and intercept) and Condition (shape or category) and their interaction in CNNs and models (D = design, B = behavioural).

| Stimulus Set | Network | Number of Layers | Model | Layer $F_{1,1}$ | Layer $p$ | Condition $F_{1,1}$ | Condition $p$ | Interaction $F_{1,1}$ | Interaction $p$ |
| --- | --- | --- | --- | --- | --- | --- | --- | --- | --- |
| A | CaffeNet | 8 | D | 92.338 | <0.001 | 14.825 | 0.002 | 4.729 | 0.050 |
|  |  |  | B | 41.233 | <0.001 | 126.651 | <0.001 | 0.202 | 0.661 |
|  | VGG-19 | 19 | D | 53.304 | <0.001 | 11.477 | 0.002 | 6.496 | 0.016 |
|  |  |  | B | 17.370 | <0.001 | 99.161 | <0.001 | 6.252 | 0.017 |
|  | GoogLeNet | 22 | D | 46.438 | <0.001 | 8.390 | 0.006 | 4.464 | 0.041 |
|  |  |  | B | 18.59 | <0.001 | 87.68 | <0.001 | 10.21 | 0.003 |
|  | ResNet50 | 52 | D | 28.788 | <0.001 | 41.075 | <0.001 | 1.662 | 0.200 |
|  |  |  | B | 25.010 | <0.001 | 750.551 | <0.001 | 0.323 | 0.571 |
| B | CaffeNet | 8 | D | 1.766 | 0.208 | 173.677 | <0.001 | 29.577 | <0.001 |
|  |  |  | B | 8.306 | 0.014 | 212.106 | <0.001 | 7.774 | 0.016 |
|  | VGG-19 | 19 | D | 2.567 | 0.118 | 160.955 | <0.001 | 4.023 | 0.053 |
|  |  |  | B | 22.075 | <0.001 | 207.91 | <0.001 | 3.536 | 0.069 |
|  | GoogLeNet | 22 | D | 2.026 | 0.162 | 312.186 | <0.001 | 8.551 | 0.006 |
|  |  |  | B | 27.727 | <0.001 | 329.938 | <0.001 | 1.833 | 0.183 |
|  | ResNet50 | 52 | D | 26.517 | <0.001 | 1377.504 | <0.001 | 0.012 | 0.913 |
|  |  |  | B | 61.007 | <0.001 | 1108.272 | <0.001 | 0.311 | 0.578 |

In summary, across both Sets A and B, we can see that shape information gradually increases and/or wavers as the network is traversed, before falling in the final layers. The peak value in shape information remains roughly the same regardless of network depth. Peak category correlations also remain roughly the same regardless of network depth. Across both Sets A and B, category information is at or below the significance threshold in the initial layer before reaching the maximum value at the final layer, showing the opposite trend with shape correlations. Interestingly, the maximum levels of shape and category correlations do not depend on network depth, nor on architectural design differences, such as the use of inception modules. Figure 3 contains multidimensional scaling plots of peak design shape and category information for Sets A and B.

**Figure 3:**
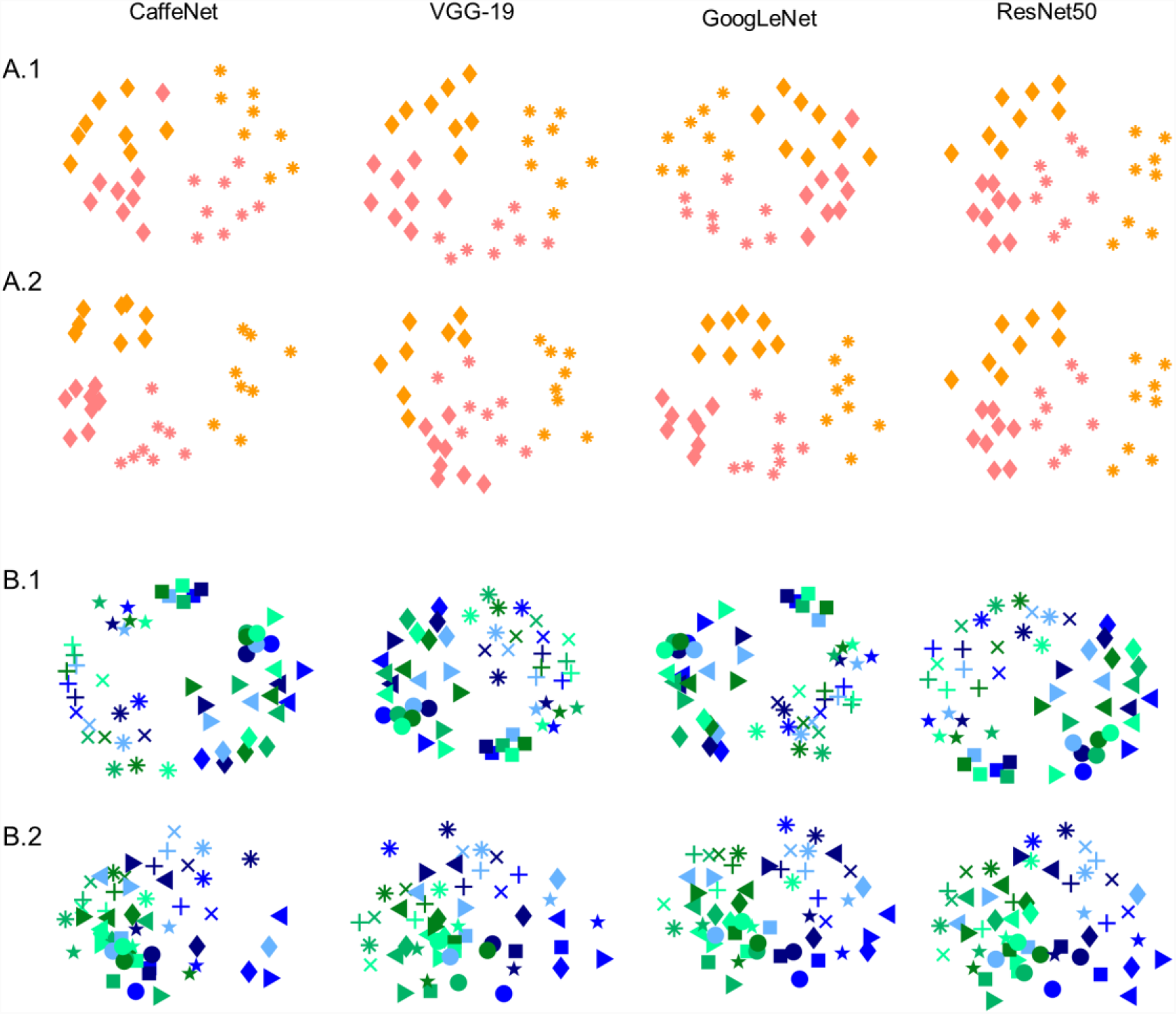
Multidimensional scaling plots of 1) Peak design shape correlations with common shape represented by common symbols, and 2) peak category correlations, with common category represented by shared colour, for each network and Set A (top 2 rows) and B (bottom 2 rows). Colour coding corresponds to Figure 1.

### Shape Versus Category information in Visual Ventral Stream Regions

Figure 4 summarises the representational similarity in three regions of interest (ROIs) along the visual ventral pathway, from low-level area V1 through to posterior and anterior VTC, compared with design and behavioural models of shape and category. For Set A, shape information reduces slightly along the ventral stream, from 22% to 19% in design models, and 18% to 10% in behavioural models. Category information increases along the ventral pathway, from −3% to 41% in design models, and −6% to 40% in behavioural models. We tested RSA results using a two-factor ANOVA, with ROI (V1, VTC ant, VTC post) and Condition (category, shape) as within-subject factors. For Set A, results reveal a significant main effect for ROI (*F*_2,15_ = 26.34, *p* < 0.001 for the design model; *F*_2,15_ = 35.81, *p* < 0.001 for behavioural), whereas the main effect of Condition (shape vs category) is not significant (*F*_1,15_ = 0.56, for design; *F*_1,15_ = 1.02, for behavioural). There is a significant interaction between ROI and Condition (*F*_2,15_ = 68.14, *p* <0.001 for design, *F*_2,15_ = 73.34, *p* < 0.001 for behavioural), indicating that as category information increases from V1 to VTC ant, shape information decreases. Post hoc pairwise t-tests further confirmed the dissociation between shape and category along the visual ventral stream: category divisions were able to significantly better explain the neural pattern in later ventral areas (VTC ant) relative to shape (*t*_(15)_ = 8.57, *p* < 0.0001 for design models, *t*_(15)_ = 5.67, *p* < 0.0001 for behavioural models); whereas the opposite was true in early visual area V1, where shape was significantly more related to the neural data compared to category divisions (*t*_(15)_ = 6.34, *p* < 0.0001 for design models, *t*_(15)_ = 8.16, *p* < 0.0001 for behavioural models).

**Figure 4:**
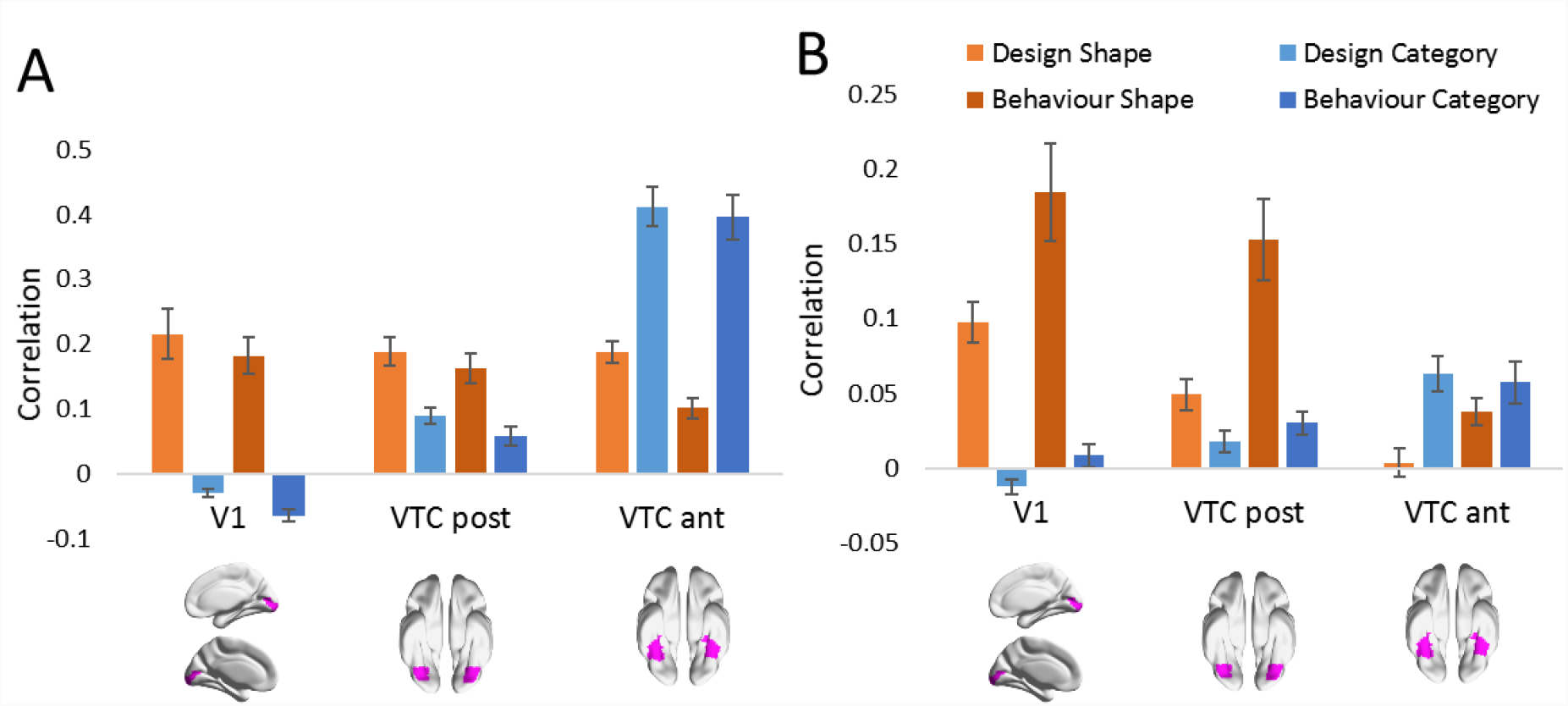
RSA results for shape and category models for Set A (left) and B (right) in ROIs. Three regions along the ventral visual pathway are analysed: V1, VTC post and VTC ant. Error bars represent standard error. ROI visualisations are re-used from Fig 4A in (Bracci, Kalfas, & Op de Beeck^33^, p. 8). Note the difference in scale between A and B.

For Set B, we see a qualitatively similar trend of decreasing shape information from V1 to VTC anterior (from 10% to 0% in the design models, and from 18% to 4% in the behavioural models) and increasing category information (from −1% to 6% in the design models, and from 1% to 6% in the behavioural models). The two-factor ANOVA, with ROI (V1, VTC ant, VTC post) and Condition (category, shape), revealed that when correlating ROI representations with the design models for Set B, ROI has no significant effect (*F*_2,14_ = 0.57, ns), the effect of Condition is significant (*F*_1,14_ = 11.39, *p* < 0.01) and there is a highly significant interaction effect between area and condition (*F*_2,14_ = 36.71, *p* < 0.001). Analysing correlations with the behavioural models for Set B, the effect of area is significant (*F*_2,14_ = 3.79, *p* = 0.027), as is condition (*F*_1,14_ = 33.84, *p* < 0.001) and there is a highly significant interaction effect between area and condition (*F*_2,14_ = 13.33, *p* < 0.001). Again, pairwise t-tests further confirmed the dissociation between shape and category in visual ventral brain regions, with shape being significantly more related to neural data in early visual area V1 than category (*t*_(14)_ = 7.56, *p* < 0.0001 for design models, *t*_(14)_ = 5.28, *p* = 0.0001 for behavioural models); and category able to explain neural patterns more in VTC ant than shape (significantly for design models *t*_(14)_ = 3.89, *p* = 0.0007, but not significantly for behavioural models: *t*_(14)_ = 1.20, *p* = 0.24). Thus, there is a two-way interaction between shape and category across the visual ventral stream that is significant for both stimulus sets and both model types, illustrating a decrease in shape combined with an increase in category going from V1 to VTC anterior.

### RSA for fMRI Brain Data and all CNN layers

Neural fMRI responses for each participant, and ROI, for Set A and Set B were correlated with the RDMs of every layer for each CNN. Results are shown in Figure 5. For each stimulus set and network, correlation values were tested in a 2 × 3 ANOVA with Layer (modelled linearly with intercept and slope) and ROI as within subject factors. In CaffeNet, V1 and VTC posterior correlations peaked at the third convolutional layer, and VTC anterior peaks at the final layer for both stimulus sets. For both stimulus sets, the 2 × 3 ANOVA results reveal a significant main effect of ROI (Set A: *F*_2,15_ = 88.73, *p* < 0.001; Set B: *F*_2,14_ = 57.00, *p* < 0.001) and Layer (Set A: *F*_1,15_ = 41.06, *p* < 0.001; *F*_1,14_ = 48.38, *p* < 0.001) and their interaction (Set A: *F*_2,15_ = 133.72, *p* < 0.001; Set B: *F*_2,14_ = 44.88, *p* < 0.001). In VGG-19, both stimulus sets show similar peaks in correlations, with V1 reaching a maximum at layer 13, VTC posterior at layer 15, and VTC anterior at the final 19^th^ layer. For both sets, there is a significant main effect of ROI (Set A: *F*_2,15_ = 59.12, *p <* 0.001; Set B: *F*_2,14_ = 26.98, *p* < 0.001) and Layer (Set A: *F*_1,15_ = 294.14, *p* < 0.001; *F*_1,14_ = 40.30, *p* < 0.001). The ROI x Layer interaction is significant in Set A (*F*_2,15_ = 55.49, *p* < 0.001), but does not reach significance in Set B (*F*_2,14_ = 2.76, *p* = 0.06). GoogLeNet has multiple peaks for correlations with V1 and VTC posterior, and there is a clear peak in VTC anterior in the final layer for both stimulus sets. For both Sets, ROI (Set A: *F*_2,15_ = 73.76, *p* < 0.001; Set B: *F*_2,14_ = 37.07, *p* < 0.001), Layer (Set A: *F*_1,15_ = 152.19, *p* < 0.001; Set B: *F*_1,14_ = 18.08, *p* < 0.001) and their interaction (Set A: *F*_2,15_ = 130.85, *p* < 0.001; Set B: *F*_2,14_ = 12.46, *p* < 0.001) are all highly significant. Finally, in ResNet50, V1 peaks at layers 44 to 47, VTC posterior peaks at layers 47 to 49, and VTC anterior peaks at the final layer. For both Sets, ROI (Set A: *F*_2,15_ = 31.20, *p* < 0.001; Set B: *F*_2,14_ = 20.26, *p* < 0.001) and Layer (Set A: *F*_1,15_ = 1431.40, *p* < 0.001; Set B: *F*_1,14_ = 895.32, *p* < 0.001) are highly significant, and their interaction is significant (Set A: *F*_2,15_ = 5.97, *p* = 0.003; Set B: *F*_2,14_ = 52.54, *p* < 0.001). Together these results show that across all deep neural networks, there is a cascade in correlation peaks from V1 to VTC posterior to VTC anterior along the layers of each network, matching with the flow of activation along the human visual ventral pathway. For all networks, and both stimulus sets, the highest correlation of VTC anterior occurs at the final layer.

**Figure 5:**
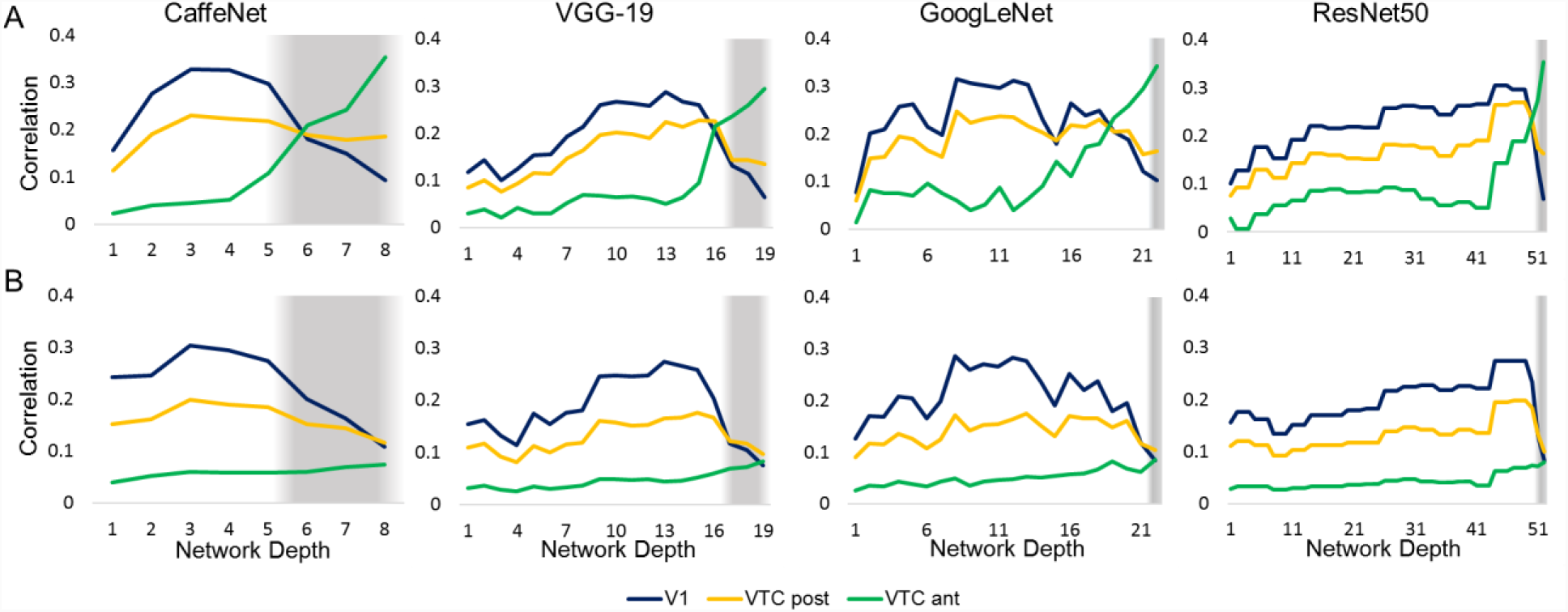
RSA comparing models (CaffeNet, VGG-19, GoogLeNet and ResNet50) and fMRI activation in V1 (navy), VTC post (yellow) and VTC ant (green) ROIs for Sets A (top row) and B (bottom row). Grey shading indicates fully-connected layers.

## Discussion

In this study, we investigated orthogonal shape and category representations in biological and artificial networks by making comparisons between: (i) CNNs and models of shape and category; (ii) models and the brain; and (iii) CNNs and the brain. First, comparing artificial networks and models, we found that CNNs represent category information as well as shape, and that category information peaks at the final layer for all tested CNNs, regardless of network depth. Peak correlation levels for shape and category do not increase with network depth, and remain roughly at the same level regardless of architectural design differences, including the use of inception modules or residual networks. Second, comparing models and the brain, there is a two-way interaction between shape and category in the human visual ventral pathway, where shape is best represented earlier in V1, and category emerges later in anterior VTC. This interaction between shape and category is significant across both stimulus sets and for both design and behavioural models. Third, comparing artificial networks and the brain, V1 correlates highest with early to mid-level layers of deep networks, and anterior VTC correlates best with the final layer of CNNs. Across both stimulus sets and for all networks, peak correlations with V1 always occur in earlier network layers than peak correlations with anterior VTC, demonstrating that CNNs reflect a similar order of computational stages as the human ventral pathway when processing these object images.

Our results allow for a greater understanding of how shape and category are represented in deep networks and in the visual ventral pathway, in particular: (i) how differing shape and category definitions between the two stimulus sets reveal differences between low-level and high-level shape representations in CNNs and the brain; (ii) how shape and category processing along deep network layers maps onto brain regions; and iii) how careful stimulus design allows us to make better inferences about category semantics in the brain and in CNNs.

One major advantage of this study is that we consider two stimulus sets that carefully control shape and category to draw conclusions about their interaction and interplay, rather than broadly extrapolating results based on a single set of images. These two well-controlled stimulus sets are similar in design but differ slightly in how shape and category are defined, allowing us to extract a finer interpretation of results. Looking at the differences in shape definitions between these stimulus sets, in Set A, shape is defined with a low to high aspect ratio (described as “bar-like” or “blob-like”), while it is characterized retinotopically in Set B. Comparing CNNs and models, both low-level (Set B) and high-level (Set A) shape information is preserved until the very last layer of all networks, however there is a visible reduction in low-level compared to high-level shape information in the final layers. Comparing models and the brain, we see that the high-level (Set A) shape information remains quite high in VTC ant, compared to low-level (Set B) shape information, which reduces to correlation levels that are at or near zero. The plausible explanation for why shape information drops off in Set B but not in A, is that higher level regions represent a more abstract form of shape, which is factored into the design of Set A, but not B. Indeed, previous studies showed that perceived shape similarity strongly overlaps with higher-level brain representations in humans^40^, and in monkeys^12,41^. Kalfas *et al.*^12^ *found that the* deepest layers of networks, rather than IT responses, correlated best with human shape similarity judgements. We also found that CNNs correlated much higher with behavioural shape judgements than fMRI. This finding suggests that there is at least some correspondence between how humans and models use shape, even though there are very likely also differences (see e.g. Baker et al.^19^).

Considering the differences in category definitions between the stimulus sets, Set A has only two category clusters defined by the animate-inanimate division, whereas Set B has six object clusters. The number of groups clearly affects the size difference in correlation levels between category models and CNNs as well as the brain, where fewer groupings boost the signal. In the final layer of all CNNs, we see that category, as defined by animacy in Set A, reaches correlation levels up to three times the magnitude of Set B. Considering brain data, category as defined by animacy in Set A reaches six times the magnitude in VTC ant compared to Set B. This is consistent with existing studies that show a strong animacy division in higher-level regions of visual cortex^24^. We find that in all four networks, human similarity judgements of category are best explained by the final layer of CNNs, more so than fMRI representations in late ventral areas.

Our use of multiple CNNs allows us to observe the influence of network depth on peak correlations with brain regions. Hong et al.^9^ compared their brain data to a CNN consisting of 6 parallelised convolutional layers, finding that the model’s top hidden layer was most predictive of IT response patterns and that lower layers had higher resemblance to V1-like Gabor patterns. Consistent with their findings, we also found that the final layer of CNNs had maximum correspondence with later ventral stream areas, and that earlier layers showed higher correlation with V1. Cichy et al.^14^ found peak V1 correlations in the second layer of an 8-layer CNN trained for object recognition. Similarly in our experiments, we found that peak V1 correlations occurred at layer 3 in an 8-layer network (CaffeNet) for both stimulus sets. As network depth increases, peak correlations with V1 shift from earlier tiers in the network to later layers. Interestingly, some of the highest V1 correlations occur immediately prior to fully connected layers, as is the case in ResNet50 and VGG-19. Figure 5 illustrates peak V1 correlations occurring as late as the 45^th^ layer in ResNet50, bringing into question the explanatory value of additional processing stages in deeper networks, especially when an 8-layer network achieves similar magnitudes of correlation with V1 by the third layer. Nevertheless, while the maximum correlation values of brain regions shift to later layers in larger networks, the rank-order of correlation peaks with brain regions still matches the order of information processing along the ventral pathway. That is, correlations with V1 always peak before VTC ant, regardless of network depth. We extend upon the findings of Cichy et al.^14^ on the order of visual information processing from a single 8 layer network to multiple networks, including a 50 layer network.

Recently, there has been some effort directed towards investigating the role of semantic representations in deep visual networks, and where category semantics may be represented in the ventral pathways^13^. Deriving high-level semantic meaning from low-level feature descriptions is commonly referred to as the “semantic gap” in computer vision literature^42^. In order to fully establish the level at which CNNs are able bridge the semantic gap, and extract meaningful information from images, it is necessary to remove all possible reliance on low-level features, which could be exploited to improve performance, and test network performance on carefully designed images that minimise potential dependencies between category and influencing features. Devereux et al.^13^ do not properly control for the influence of shape, as we have, and include many low-level visual features labelled misleadingly as “semantic” descriptors, such as “is circular/round” or “is “green”, which we would argue do not allow for a dissociation between vision and semantics^15^. Our study explicitly defines category semantics as falling within the animacy division in Set A, or in multiple object categories (animals, minerals, fruit/vegetables, music, sports equipment and tools) in Set B. Our stimulus sets do not confound category semantics with shape information, allowing us to draw firmer conclusions.

In conclusion, despite shape and category often being confounded in natural images, and the possibility for artificial neural networks to exploit this correlation when performing classification tasks, we find that deep convolutional neural networks are able to represent category information independently from low-level shape in a manner similar to higher level visual cortex in humans.

## Acknowledgements

A.A.Z. and H.O.d.B were funded by grant C14/16/031 of the KULeuven Research Council. J.B.R. received funding from the FWO and European Union’s Horizon 2020 research and innovation programme under the Marie Skłodowska-Curie grant agreement No 665501, via a FWO [PEGASUS]^2^ Marie Skłodowska-Curie fellowship (12T9217N). S.B. is funded by FWO (Fonds Wetenschappelijk Onderzoek) postdoctoral fellowship 516 (12S1317N). Neuroimaging was funded by the Flemish Government Hercules Grant ZW11_10. We would like to thank Tim Leers for help with data analysis.

## Data Availability

The datasets generated during and/or analysed during the current study are available from the corresponding author on reasonable request.

## Author Information

### Contributions

All authors contributed to the study design. SB, JBR and HOdB provided pre-processed neuroimaging data and collected behavioural data. AAZ ran network simulations, analysed the data and wrote the manuscript with input from all authors. All authors interpreted the data, edited the manuscript and approved the final version.

